## Supplementary figures and images for "A protease protection assay for the detection of internalized alpha-synuclein pre-formed fibrils"

### raw gel and blot data

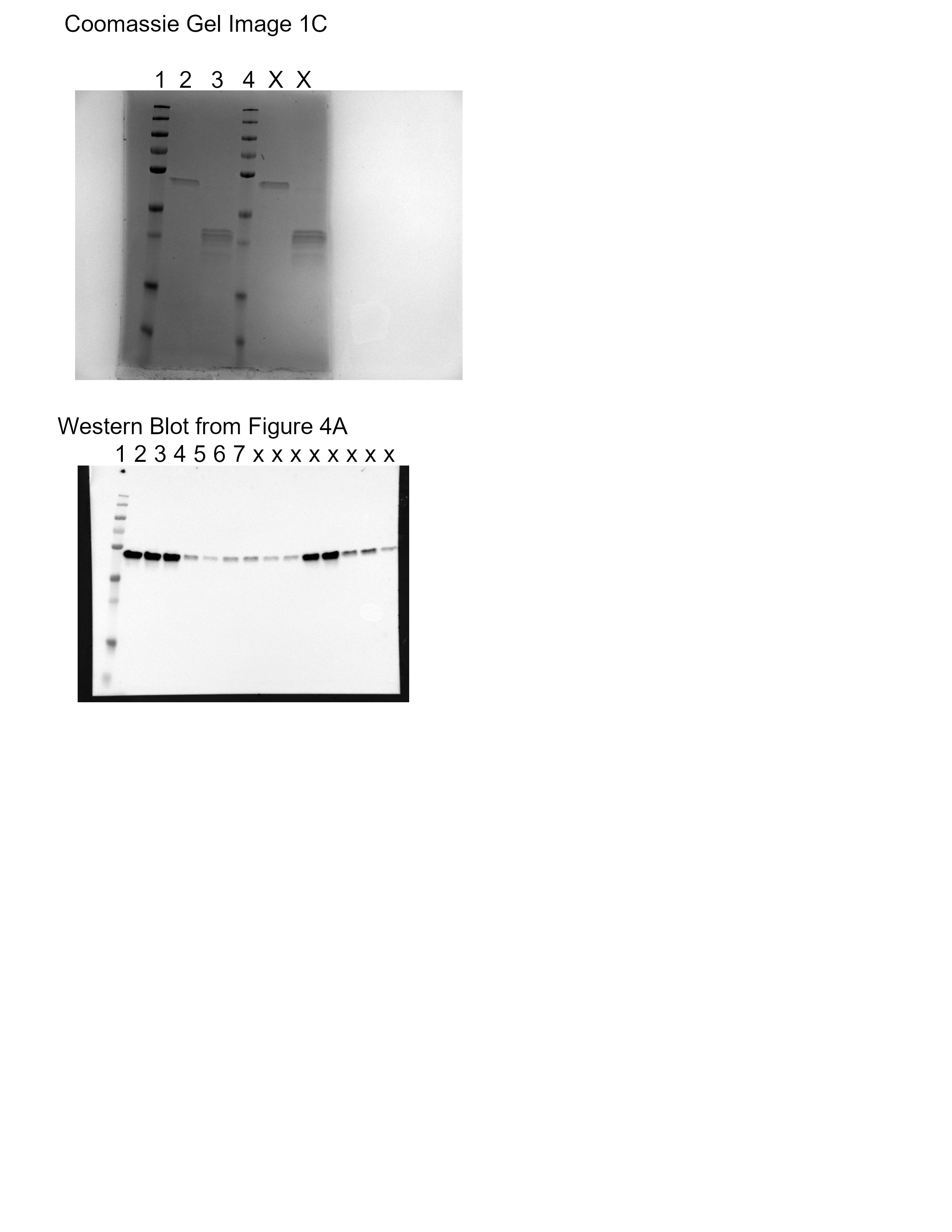
